# Cell division can accelerate the loss of a heteroplasmic mitochondrial DNA mutation in a mouse model of mitochondrial disease

**DOI:** 10.1101/2023.04.06.535821

**Authors:** Tianhong Su, Tiago M Bernardino Gomes, Anna L M Smith, Julia C Whitehall, Alasdair P Blain, Marie-Lune Simard, Louisa Scholten, James B Stewart, Doug M Turnbull, Conor Lawless, Laura C Greaves

## Abstract

Mitochondrial DNA (mtDNA) mutations accumulate in both mitotic and post-mitotic somatic tissues of normal individuals with age. They clonally expand within individual cells and cause mitochondrial dysfunction. In contrast, in patients with inherited disease-causing mtDNA mutations the mutation load decreases in mitotic tissues over time, whereas the mutations load in post-mitotic tissues remains relatively stable. The mechanisms underlying this decrease in mitotic tissues, and whether mitochondrial function is restored at the tissue level are unknown. Here, using a combination of homogenate tissue and single crypt/muscle fibre pyrosequencing we have shown a decrease in the mutation load of the germline heteroplasmic m.5024C>T mutation in multiple mitotic tissues of a mouse model of inherited mitochondrial disease (C5024T mice). We have then used *in silico* predictions to model the cellular dynamics of mtDNA mutation load in mitotic and post mitotic tissues. We demonstrate that: (1) the rate of m.5024C>T decrease correlates with the rate of tissue turnover; (2) the mutation load decrease is not associated with changes in overall cellular proliferation and apoptosis within the mitotic colonic epithelium; instead, it could be due to an upper limit of m.5024C>T load in stem cell populations; (3) the m.5024C>T mutation load is maintained in post-mitotic tissues over time with a consistent load amongst individual muscle fibres; (4) *in silico* modelling supports a scenario where genetic drift is accelerated in mitotic tissues by high levels of mtDNA replication coupled with mtDNA segregation at cell division. This study has advanced our understanding of the dynamics of mtDNA mutations and phenotype development in patients with mtDNA disease.

**Author Summary:** Healthy individuals randomly accumulate pathogenic mtDNA mutations with age in dividing cells, causing mitochondrial dysfunction. Interestingly, patients with mitochondrial disease show a relative decrease in the loads of inherited mtDNA mutations in some dividing cells over time. The mechanisms underlying this decrease are unknown. Here we show a decrease in the load of the germline heteroplasmic m.5024C>T mutation in dividing cells and tissues of a mouse model of mitochondrial disease. In contrast, the mutation load in non-dividing cells and tissue remains stable. Our data are consistent with the hypothesis that a higher frequency of mtDNA replication in dividing cells, coupled with stem cells having an upper tolerance limit for m.5024C>T, causes an overall decrease in m.5024C>T load at the tissue level.

## Introduction

Mitochondria are dynamic organelles present in most eukaryotic cells. Their primary function is the production of cellular energy in the form of adenosine 5’-triphosphate (ATP) through the process of oxidative phosphorylation (OXPHOS). Mitochondria contain their own genome, mitochondrial DNA (mtDNA), which encodes 13 polypeptides which are essential components of the OXPHOS machinery, as well as 22 tRNAs and 2 rRNAs, which are required for mitochondrial translation. Multiple copies of mtDNA are present in individual cells. When all copies of mtDNA in a cell are the same, the cell is said to be in a state of homoplasmy; however, when mutated and wild-type mtDNA co-exist in the same cell it is said to be in a state of heteroplasmy. The proportion of mutated mtDNA represents its heteroplasmy level or mutation load.

The vast majority of pathogenic mtDNA mutations are functionally recessive meaning that their mutation load has to exceed a certain threshold (typically between 60 and 95%) before a defect in OXPHOS occurs [1].

Individual human cells accumulate somatic mtDNA mutations, to high levels in some cases, in a variety of mitotic and post-mitotic tissues during ageing. These mtDNA mutations can lead to OXPHOS defects and cellular dysfunction in these tissues [2–7]. The process by which one mutated mtDNA species becomes dominant within a cell is termed clonal expansion. Mathematical modelling studies have suggested that the initial mutational events occur early in life and then mutations clonally expand over time through random genetic drift and segregation during cell division [8–10].

Germline mutations in mtDNA are also an important cause of mitochondrial dysfunction, with one in 5,000 individuals estimated to be affected by inherited mitochondrial disease [11]. Over half of the disease-causing mutations are found in tRNA encoding genes, and most of them are heteroplasmic. This results in most tissues displaying a mosaic pattern where only cells that have randomly accumulated high mutation loads are OXPHOS deficient. One of the most common mtDNA point mutations is m.3243A>G in the *MT-TL1* gene. An intriguing feature of this mutation, and many other inherited heteroplasmic mtDNA mutations, is that, in contrast to the accumulation of somatic mtDNA mutations in mitotic cells with age, the inherited mutation loads decreases over time in tissues which are rapidly dividing, such as blood, buccal mucosa, intestinal epithelium, cervical smears and epithelial cells in urine [12–16]. In contrast, the homogenate mutation load of m.3243A>G remains relatively stable in post-mitotic tissues such as heart and skeletal muscle over time [13, 14, 17–19]. Whilst selection against high loads of m.3243A>G mutation in lymphocytes (predominantly the T-cell pool) has been documented at single-cell resolution, it is not currently understood why average mutation load falls [20] in other dividing tissues, such as in epithelial cell populations, while remaining relatively unaffected in post-mitotic tissues. In addition, it is unknown whether the fall in mutation load results in a rescue of the biochemical phenotype at the tissue level.

To address these questions, we used a mouse harbouring a germline heteroplasmic m.5024C>T mutation in the mitochondrial tRNA^Ala^ gene (from here on referred to as C5024T mice) [21]. These mice display many of the genetic features of human mtDNA disease, such as germline transmission, genotype/phenotype correlations, a compensatory increase in copy number in response to mitochondrial dysfunction, and loss of the mutation in rapidly dividing blood cell lineages and intestinal epithelium over time [22]. Using a combination of clonal unit genetic analyses, mathematical modelling, and biochemical investigations we show that: 1) there is an upper limit of m.5024C>T mutation load tolerated by mitotic cells, 2) there is much less variation in individual units of post-mitotic tissues than rapidly replicating tissues, 3) the higher the rate of cell division, the more rapidly mtDNA mutations are lost.

## Results

### The mutation load of m.5024C>T decreases with age in mitotic tissues

It has been previously shown that there is a decrease in m.5024C>T mutation load in blood and intestines of C5024T mice with age [21, 22]. We wanted to confirm that this was also the case in our mouse cohort and to examine age-related m.5024C>T mutation load changes in other organs. To achieve this, we used pyrosequencing to quantify mutation loads in several tissues from 10-week [n=7 for all tissues, except pancreas (n=6), lung (n=6) and pyloric epithelium (n=3)] and 50-week C5024T mice [(n=7 for all tissues, except small intestinal smooth muscle (n=6), small intestinal epithelium (n=5), fundic epithelium (n=6) and pyloric epithelium (n=3)]. C5024T mice harboured intermediate to high mutation loads (60% - 80%) in early life, measured from ear notch obtained at weaning, and were heteroplasmy-matched between the age groups. To improve the stringency of the comparison between mice of different ages, mutation loads of each tissue were normalised by subtracting the initial mutation load quantified in the ear notch for each mouse. The difference in the normalised mutation load between 10-week and 50-week C5024T mice was calculated, revealing a significant loss of the mutation in the spleen, the epithelium of the small intestine and the gastric pyloric epithelium, with age (p=0.014, p=0.0008 and p=0.012 respectively, unpaired, two-tailed t-test, Fig. 1). In contrast, the mutation load remained the same in the intestinal smooth muscle, skeletal muscle, heart and brain, and organs that are comprised of a mix of mitotic and post-mitotic tissues and/or are mitotically quiescent under physiological conditions, such as liver, lung, and pancreas (Fig. 1). The mutation load was significantly higher in the kidney (p=0.02) and gastric smooth muscle (p=0.04, unpaired two-tailed t-test) of 50-week C5024T mice compared with 10-week C5024T mice. Surprisingly, there was no change in the mutation load with age in the mitotic epithelium of the gastric fundus.

**Figure 1:**
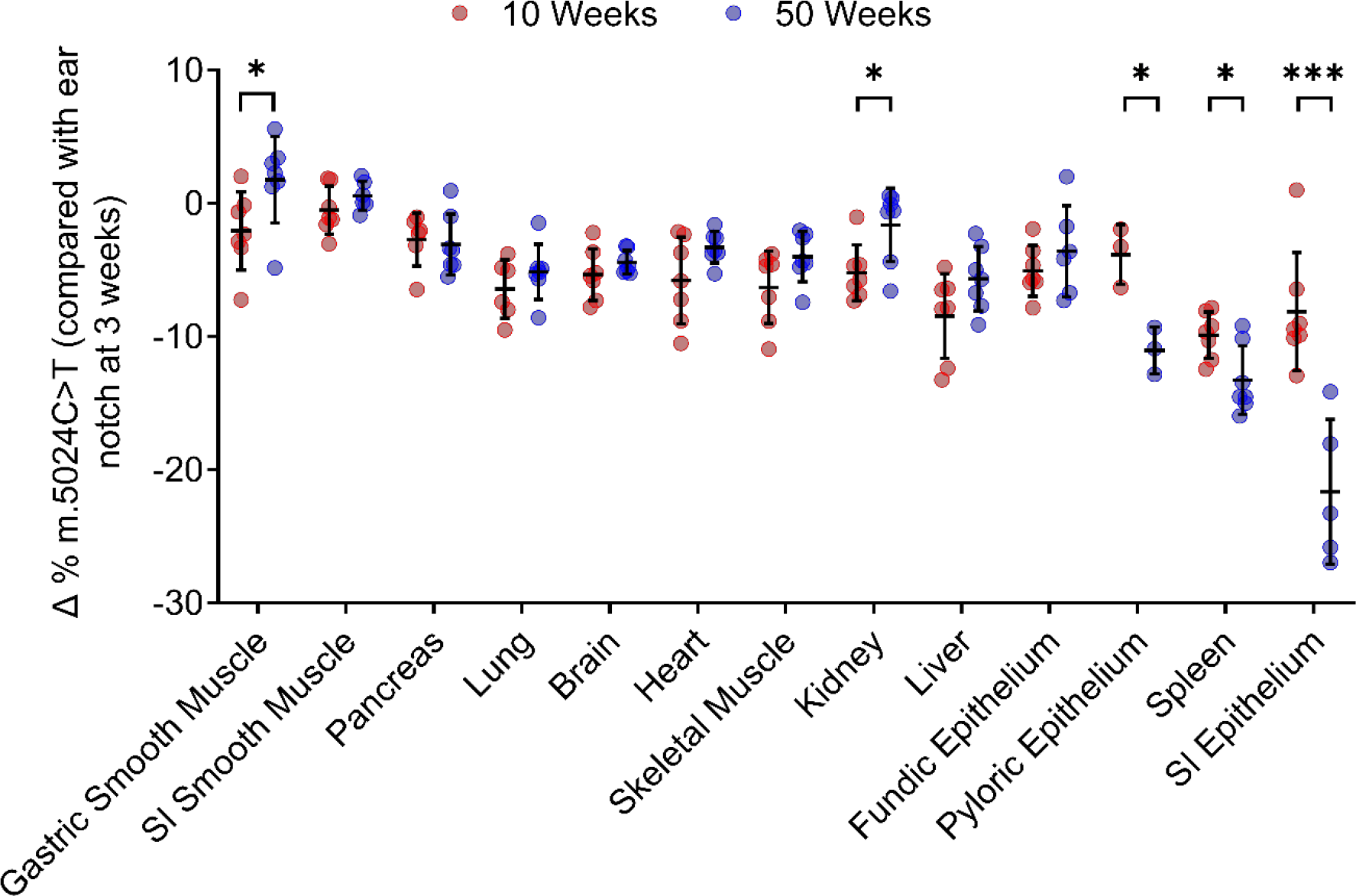
m.5024C>T mutation load in tissue homogenates from 10 and 50-week-old C504T mice compared to ear-notch mutation load at weaning (3 weeks). SI, small intestine. Lines show mean ± SD. * p< 0.05, ** p<0.01, *** p<0.001, (unpaired, two-tailed t- test)

### Mutation loss only occurs in tissues with the highest proliferation rates

The data from homogenate tissues highlighted a subset of tissues that had the highest rate of mutation loss. Tissues with the highest turnover rate (e.g., the small intestinal epithelium) had the highest rate of loss. To investigate this further, we quantified C5024T loads in two histologically and functionally distinct epithelial subtypes of the stomach; the fundic and pyloric epithelium [23]. Pyloric epithelium has a faster turnover than fundic epithelium, but both proliferate at a much lower rate than intestinal epithelium [24]. We determined the load of the m.5024C>T mutation in individual fundic and pyloric units of 50-week C5024T mice and compared that with the smooth muscle lining the stomach, which has a consistent load of m.5024C>T over time. The mutation load in pyloric units was significantly lower than that in the smooth muscle with a significantly wider variance around the mean (p < 0.0001, Kruskal-Wallis test, Fig. 2A). There was no significant difference between the mean mutation loads in fundic units compared with the smooth muscle; however, there was a significantly greater variance around the mean of the pyloric units (p < 0.01, Kruskal-Wallis test,). This suggests that there is also drift of the mutation in individual fundic units, but that it is significantly slower than in pyloric units (Fig. 2A), resulting in a slower loss of the mutation from the tissue. As a comparison, we performed a similar analysis in post-mitotic quadriceps skeletal muscle tissue and quantified m.5024C>T load in single muscle fibres from 10-week (n=6) and 50-week C5024T (n=7) mice. We found no significant difference in the mean m.5024C>T mutation load or the inter-fibre variation between 10-week and 50-week mice (p=0.65, unpaired, two-tailed t-test, Fig. 2B). Our observations suggest that mtDNA genetic drift is accelerated in dividing tissues compared with post-mitotic tissues and that this is driven by a combination of increased mtDNA replication rate and random segregation of mtDNA genotypes during cell division [25].

**Figure 2:**
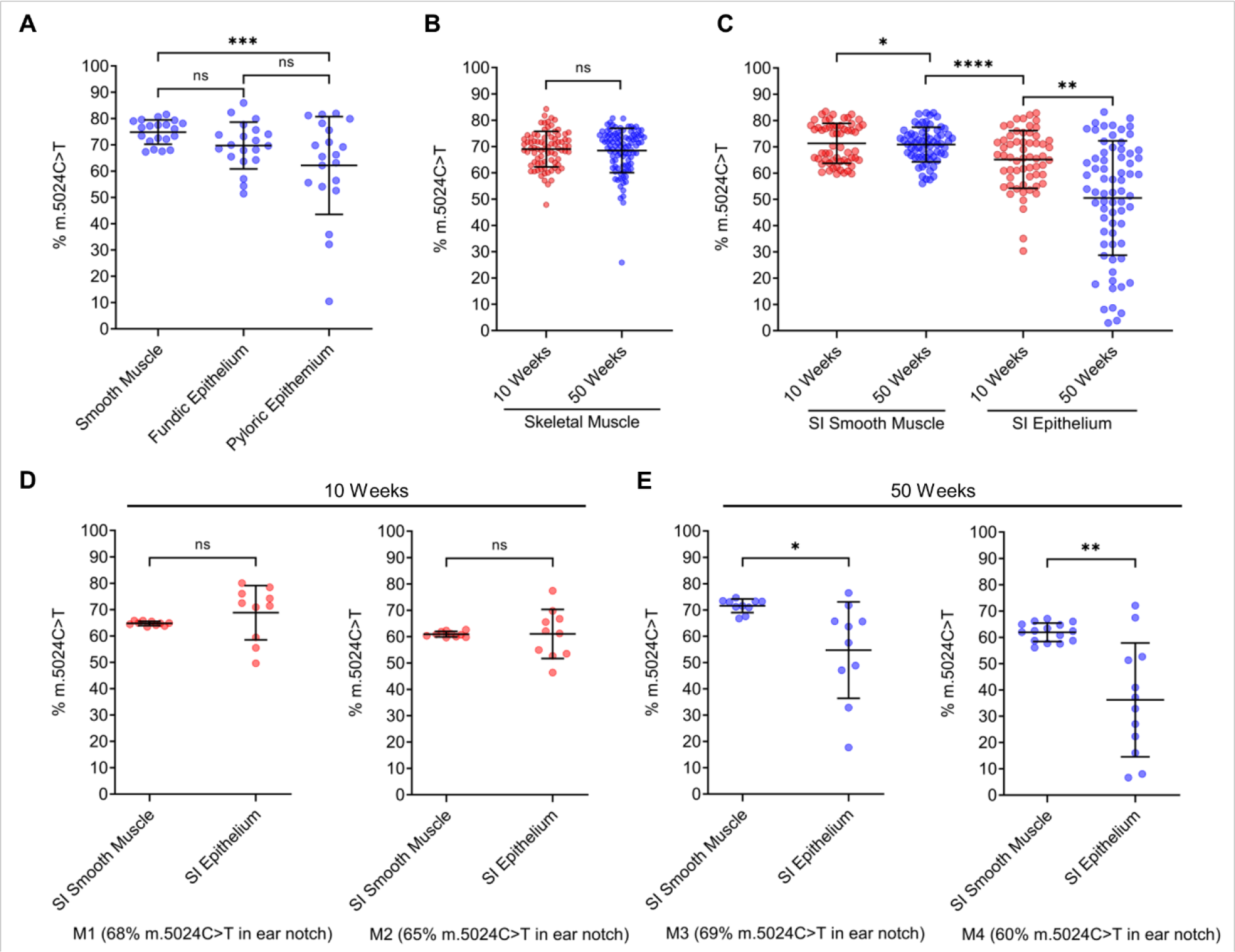
Comparison of m.5024C>T mutation loads in stomach, skeletal muscle, and small intestinal epithelium and smooth muscle. (**A**) m.5024T>C mutation load in individual gastric units of fundic/pyloric epithelium of 50-week C5024T mice measured using pyrosequencing. Each dot represents one small patch of smooth muscle or one gastric unit. Data from n=3 mice are pooled. (**B**) m.5024T>C mutation load in individual skeletal muscle fibres of C5024T mice at 10 and 50 weeks. 88 fibres were analysed from n=6 10-week mice and 100 fibres were analysed from n=7 50-week mice. Each dot denotes a fibre. (**C**) Mutation load in individual small intestinal (SI) crypts compared with that in smooth muscle patches of C5024T mice at 10 weeks (n=6 mice, n=10 patches/crypts per mouse) and 50 weeks (n=6 mice, n=10 patches/crypts per mouse). (**D-E**) Mutation levels in individual SI crypts and smooth muscle areas of two exemplar 10-week (**D**, M1-2) and two exemplar 50- week mice (**E**, M3-4) with intermediate mutation loads. Lines show mean ± SD. * p<0.05; ** p< 0.01; **** p<0.0001. (A and C)Kruskal-Wallis test; (B) unpaired two-tailed t-test; (C-D) unpaired two-tailed t-test with Welch’s correction.

### Intestinal crypts have an upper limit for tolerance of m.5024C>T

The small intestinal epithelium has the highest rate of turnover [26, 27] and the fastest rate of m.5024C>T loss of the tissues studied. Therefore, we decided to investigate mutation dynamics by comparing the distributions of mutations in individual crypts in 10-week and 50-week mice. As a comparator, we also quantified the load of m.5024C>T in the smooth muscle of the small intestine as this stays constant over time (Fig. 1). Single intestinal crypts and small patches of smooth muscle were randomly laser micro-dissected from 10-week (n = 6) and 50-week (n = 6) mice and the m.5024C>T load was quantified by pyrosequencing. The mean m.5024C>T load in intestinal crypts was significantly lower than that in smooth muscle at both 10 and 50 weeks (p<0.05 and p<0.0001 respectively, Kruskal-Wallis test), but the difference was greater at 50 weeks (Fig. 2C). Similarly, the variance was significantly higher in epithelial cells than in smooth muscle cells at both 10 weeks and 50 weeks. Further, the mean mutation load in epithelial cells was significantly lower (p<0.01, Kruskal-Wallis test, Fig. 2C) and the variance significantly higher at 50 weeks than at 10 weeks, with some crypts carrying extremely low mutation loads (∼3%). Examples of individual mice at 10 and 50 weeks are shown in Fig. 2D and 2E. Homoplasmy of m.5024C>T was never observed, instead there appeared to be an upper limit of m.5024C>T in the intestinal crypts (∼83%). All these observations are consistent with the hypothesis that mutations are lost due to random drift during replication, with selection at the cell level against high mutation loads. This process occurs at a faster rate in cells with a higher turnover resulting in the observed overall decline in the mean m.5024C>T mutation load in the intestinal epithelium (Fig. 2C).

### Loss of m.5024C>T is not associated with apoptosis or loss of cellular proliferative capacity

Having observed that intestinal crypts have a m.5024C>T load upper limit that crypt cells can tolerate before being lost, we hypothesised that this loss may be associated with a decline in replicative capacity or an increase in apoptosis of these cells. To investigate this, we performed immunofluorescence and quantified the number of proliferating cells (Ki-67), cell nuclei (Hoechst), and apoptotic cells (cleaved caspase- 3) in intestinal crypts of wild-type and C5024T mice aged 10 and 50 weeks. There was no significant difference in the proportion of proliferating cells per crypt or the frequency of apoptosis between C5024T and wild-type mice in either age group (Fig. 3). This suggests that the selective loss of cells with high mutation loads is not linked to increased apoptosis or a loss of cell proliferation capability, and is supportive of a model of stem cell loss by differentiation [22].

**Figure 3:**
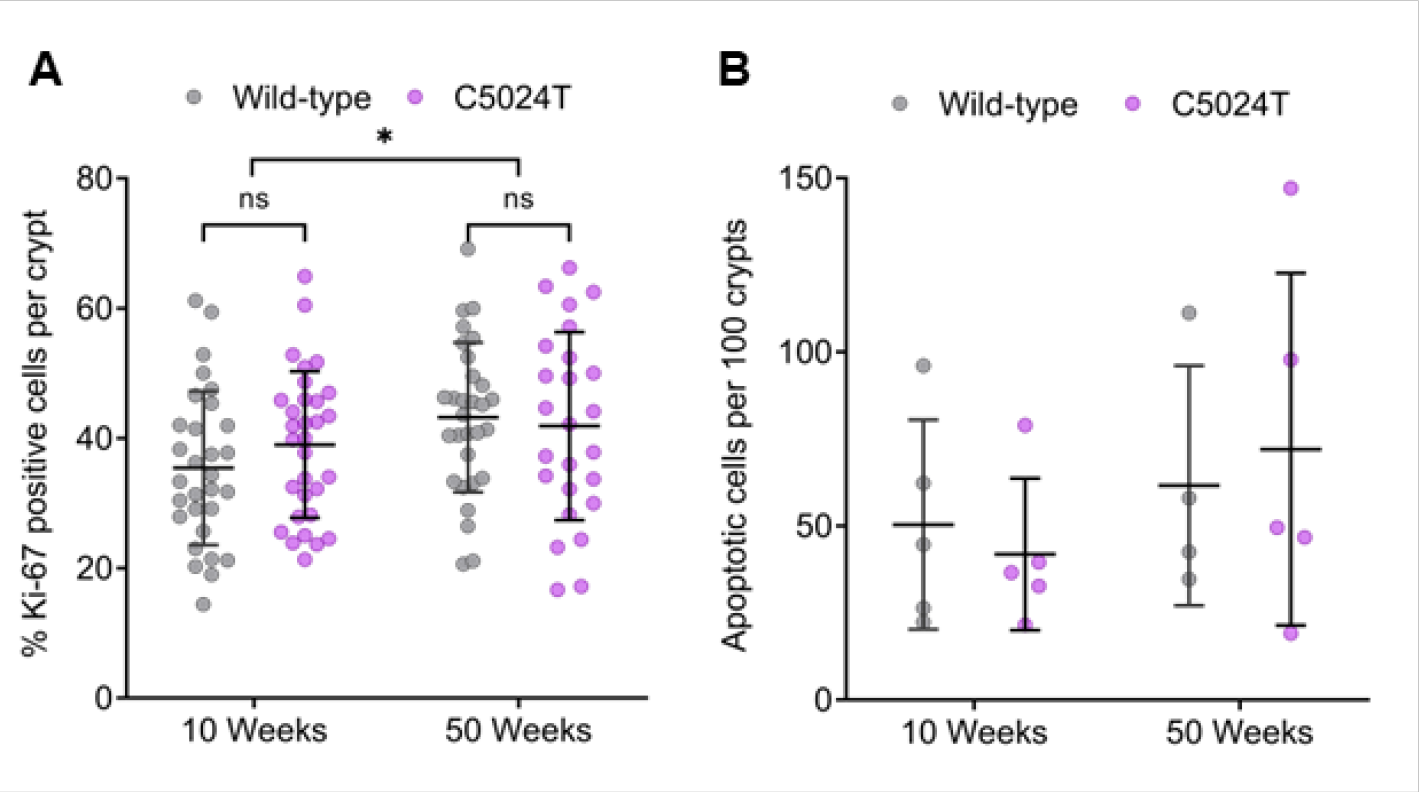
Cell proliferation and apoptosis in colonic crypts of C5024T and wild-type mice. (**A**) Percentage of Ki-67 positive proliferating cells in wild-type and C5024T mice at 10 and 50 weeks (n=10 crypts were analysed per mouse in n=3 mice per group). Each dot represents an individual crypt. (**B**) Apoptosis levels were measured in 10-week and 50-week C5024T mice vs and compared with wild-type mice. The frequency of apoptotic cells per 100 crypts per mouse was quantified. Each symbol signifies a mouse (n=5 C5024T and n=4 wild-type mice per age group). All results were analysed using two-way ANOVA and presented as mean ± SEM, *p<0.05.

### Stringent mtDNA replication can explain the difference between mutation dynamics in skeletal muscle fibres and intestinal epithelial crypts

To test the hypothesis that cell division accelerates mutation loss, we developed a mathematical model to describe the replication of mtDNA and expansion of mtDNA mutations in two contrasting types of tissue: (1) non-dividing cells (e.g., post-mitotic skeletal muscle fibres) and (2) stem cells dividing in an epithelial stem cell niche. Our model simulates random genetic drift due to the usual relaxed mtDNA replication, independent of the cell cycle [8, 10], as well as random drift due to stringent mtDNA replication, linked to the cell cycle [28], uneven segregation of mtDNA molecules during cell division and a loss of stem cells (via loss of replicative competitiveness within its niche) once mutation load exceeds a threshold. We used the model to simulate mtDNA dynamics in skeletal muscle fibres and intestinal epithelial stem cell niches in C5024T mice and compared the results with experimental observations (Fig. 2 and Fig. S1). Model parameters were based on published experimental data wherever possible (Table 1).

**Table 1:** Simulation parameters used in the in silico modelling for non-dividing cells and stem cell populations.

| Parameters common to both models | Semantic explanation | Parameters specific to the stem cell niche simulation | Semantic explanation |
| --- | --- | --- | --- |
| $r_w = 0.001875 d^{-1}$ | The rate constant describing replication of wild-type molecules according to first order mass-action kinetics | $N_{targ} = 7 [30]$ | The target, steady-state number of stem cells |
| $d_w = 0.0075 d^{-1}$ | The equivalent degradation rate to above | $\mu = 3 d [30]$ | Mean time between divisions |
| $r_m = 0.001875 d^{-1}$ | The rate constant describing replication of mutant molecules | $\sigma = 0.1 d$ | SD of the above |
| $d_m = 0.0075 d^{-1}$ | The degradation rate of mutant molecules | $\delta = 2$ | Difference between the number of stem cells in a niche and the target, steady-state number of stem cells |
| $m = 0.00003 [51]$ | The probability that a replication event gives rise to a mutant mtDNA molecule | $P = 0.75 [10]$ | The probability of asymmetric divisions |
| $T = 200 [52]$ | The steady-state total mtDNA population size | | |
| $L = 0.875$ | The upper limit above which the mtDNA population is lost | | |

#### Modelling the effect of relaxed replication on mtDNA population dynamics

This model describes a population of wild-type and mutant mtDNA molecules undergoing relaxed replication within a single cell. The initial mutation load distribution was estimated from experimental observations of mutation loads from individual skeletal muscle fibres of 10-week mice. The population evolves, simulating each mtDNA replication and degradation event in a cell until a defined endpoint (e.g., approximately the lifespan of the mouse) or until the cell mutation load exceeds a previously determined lethal limit. When the mutation load exceeds the upper limit, the cell and its entire mtDNA population is assumed to be lost. mtDNA replication and degradation rates are balanced to maintain approximate mtDNA copy number homeostasis.

#### The additional effect of stringent replication on mtDNA population dynamics in the intestinal epithelium stem cell niche

To examine the additional effect of stringent replication during cell division, we expanded the model to also simulate mtDNA dynamics within the intestinal epithelial crypt stem cell niche. We assume the same initial mutation load distribution, as described above, and that the number of ‘effective’ stem cells in the niche is approximately 7 [29, 30]. Each stem cell divides asynchronously approximately every three days [30] and the mtDNA population doubles prior to cell division before randomly segregating into the two daughter cells. Stem cell division symmetry is assigned randomly, with asymmetric division occurring with probability 0.75. Symmetric division with stem-cell fate and symmetric division with non-stem cell fate occur with equal probability of 0.125, maintaining approximate homeostasis in niche size. However, if the number of stem cells within the niche drops below 5, then division is always symmetric with stem-cell fate. If the number of stem cells within the niche exceeds 9, then division is always symmetric with non-stem cell fate [29, 30]. mtDNA dynamics between cell divisions were modelled using the relaxed replication model above, with the same parameters.

#### Comparing simulated mtDNA dynamics with experimental observations from C5024T mice

Using the model above, we simulated mtDNA population dynamics with stringent and relaxed replication to represent mitotic epithelial crypts, and only relaxed replication for post-mitotic skeletal muscle. We ran 1,200 simulations of stem cell niches representing ∼8,400 crypt stem cells and 12,000 simulations representing 12,000 theoretical individual muscle fibres. Both sets of simulations were carried out until animals were 100 weeks old. The relaxed replication rate parameter was chosen as the highest value that gave simulation results that fit both sets of data by visual comparison between the simulated mutation load distributions and their corresponding experimental observations of single epithelial crypts or skeletal muscle fibres from both 10-week and 50-week C5024T mice.

Our simulations show that there exist model parameters consistent with a negligible change in the m.5024C>T mutation load distribution in skeletal muscle fibres over time, and a significant widening of this distribution in intestinal crypts, resulting in a decrease in average mutation load across the lifespan of the mice. The simulated data are in close agreement with our experimental data (Fig. 4). In skeletal muscle, the experimentally observed m.5024C>T mutation loads in both 10-week and 50-week mice were tightly distributed with means of 69% (SD=6.7%) and 68% (SD=8.4%). The equivalent simulated results are similar (Fig. 4A), with means of 74% (SD=5.6%) and 73% (SD=7.6%). However, in epithelial crypts, experimentally observed m.5024C>T mutation loads in 10-week mice have a mean of 65% (SD=11%) and in 50-week mice 51% (SD=21.7%). The equivalent simulated results show a means of 71% (SD=6.6%) and 60% (SD=15.2%), respectively (Fig. 4B). These data support our hypothesis that random genetic drift and an upper limit for tolerance of m.5024C>T, coupled with high rates of cell division can explain the significant decline in the m.5024C>T mutation in dividing cells with age without requiring selective removal mechanisms such as apoptosis.

**Figure 4:**
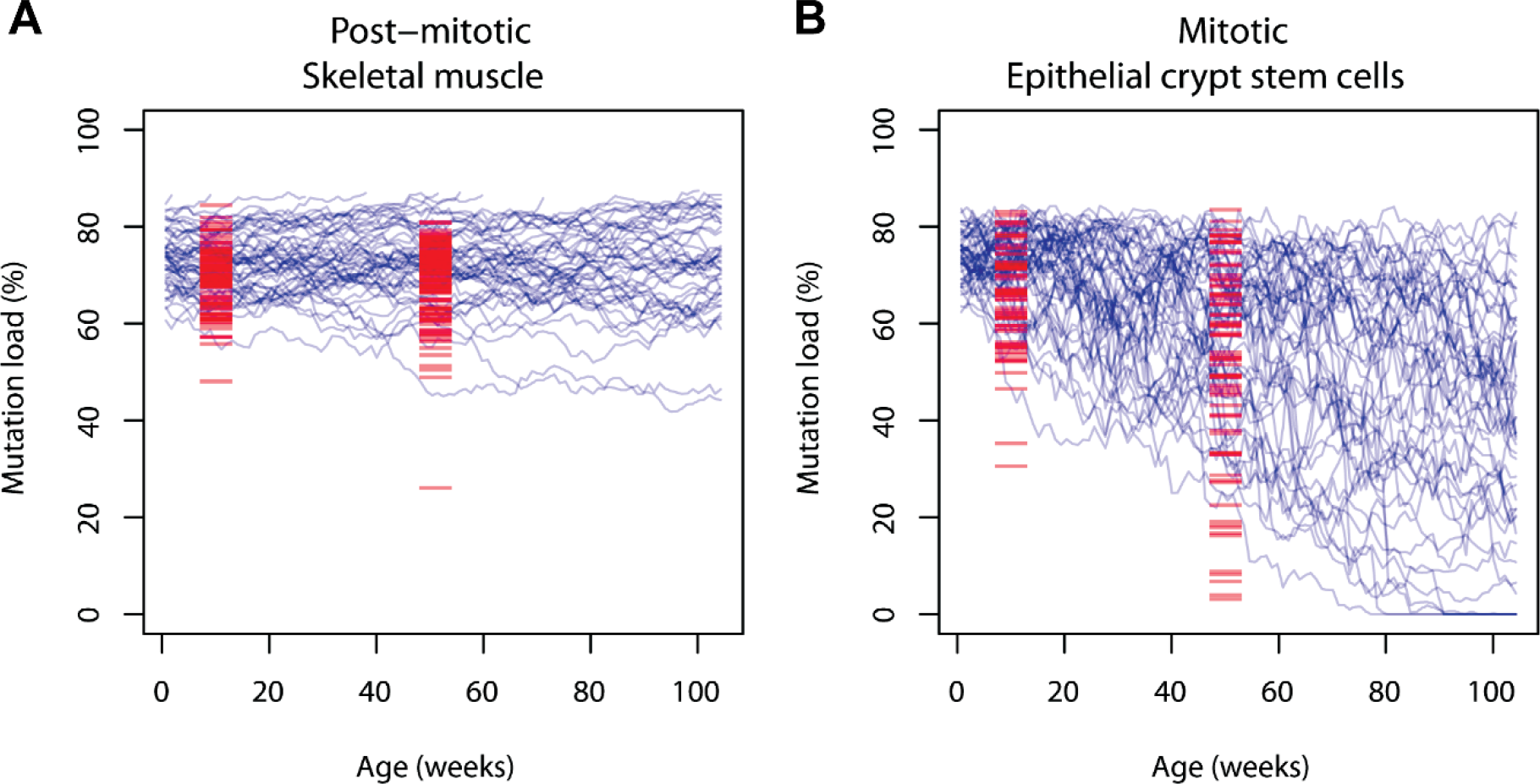
**A comparison of experimental observations of mtDNA population dynamics with model predictions**. (**A**) 88 experimental observations of mutation load in skeletal muscle fibres from 10-week mice and 100 observations from 50-week mice (red) together with 150 simulations of mutation load dynamics drawn randomly from 1,200 simulated fibres across 100 weeks (blue). (**B**) 60 experimental observations of mutation load in intestinal epithelial crypts from 10-week mice and 69 observations from 50-week mice (red) together with 150 simulations of mutation load dynamics drawn randomly from 1,200 simulated crypt stem cells (blue).

### A decrease in m.5024C>T mutation load may not rescue the OXPHOS phenotype in intestinal epithelial crypts

Next, we wanted to quantify OXPHOS function in the crypts of the C5024T mice to determine whether the decline in the mutation load improved OXPHOS function relative to age-matched control wild-type mice. To this end, we used a validated histochemical protocol, NBTx [31], to quantify the activity of cytochrome c oxidase (complex IV) in individual colonic crypts. NBTx enables precise quantification of complex IV enzyme activity by using competing redox reactions which ultimately lead to the reduction of tetrazolium salts and formazan deposition in cytochrome *c* oxidase (COX)-deficient mitochondria (Fig. 5A). The levels of this reaction product were quantified using densitometry in a minimum of 100 individual colonic crypts of C5024T and wild-type mice of 10 and 50 weeks of age. In 10-week mice, an average of 19% (SD±11.53) of crypts were COX-deficient in the C5024T mutants (Fig. 5B-D). This correlated significantly with the initial ear notch mutation load at 3 weeks (p=0.03, Spearman’s Rank Correlation, Fig. S1A). None of the crypts in the 10-week wild-type mice were COX-deficient (Fig. 5). Interestingly, despite the significantly lower mutational load observed in 50-week C5024T mice, when compared to 10-week mutants, there was no significant change in the proportion of COX-deficient crypts (p=0.41, Mann-Whitney Test), though the numbers of mice available for analysis were low. For 50-week C5024T mice, there was no correlation between the proportion of COX-deficient crypts and the initial m.5024C>T mutation load (Fig. S1B) suggesting other mechanisms may be contributing to preserving the established COX deficient phenotype in these mice.

**Figure 5:**
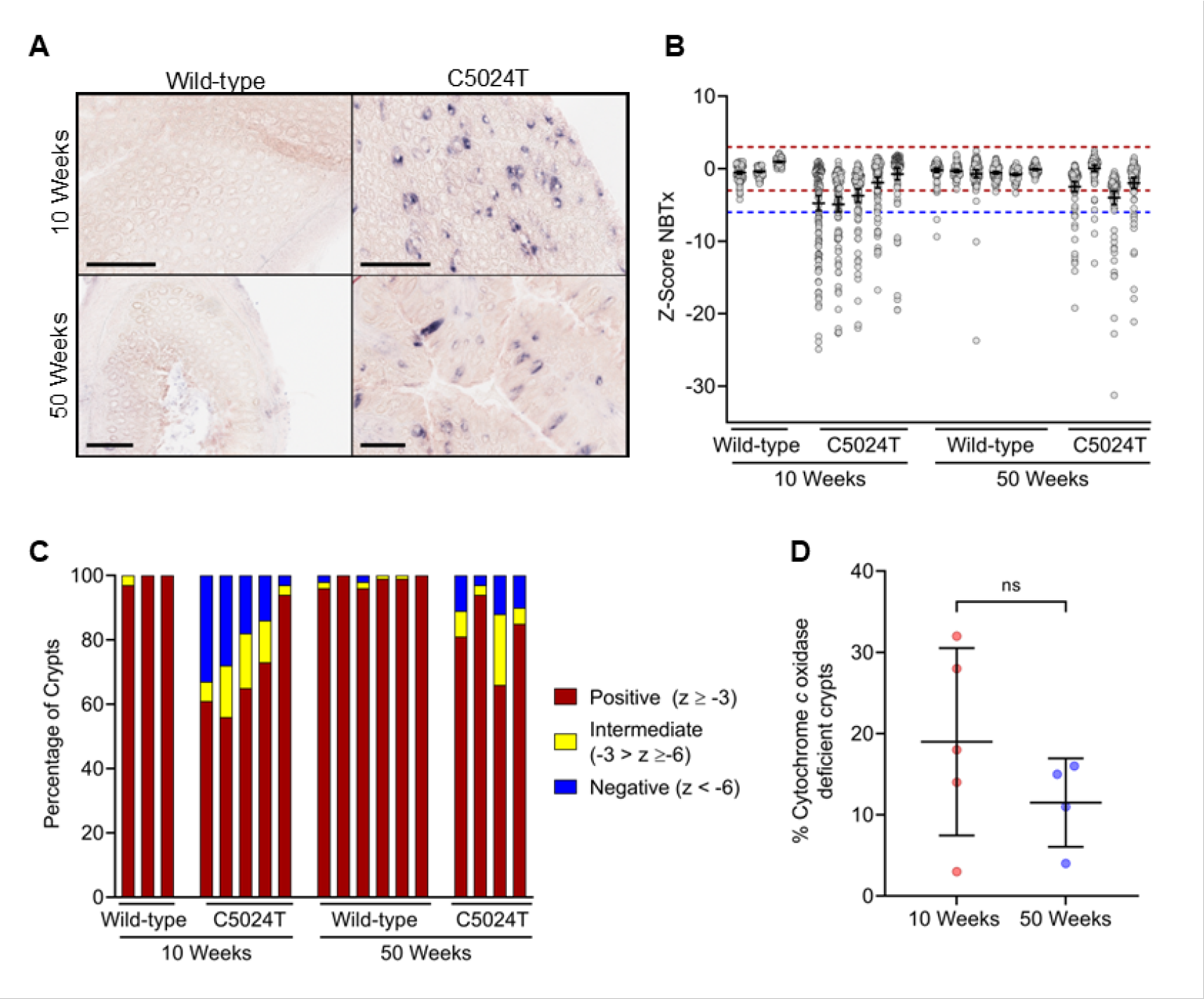
Analysis of Cytochrome *c* Oxidase (COX) activity in colonic crypts of C5024T and wild-type mice with 10 and 50 weeks of age. (**A**) Representative images of NBTx histochemical staining of colonic crypts to demonstrate COX activity (scale bars 200 µm). The blue histochemical reaction product shows loss of normal COX activity, whereas a lack of reaction product shows normal COX activity. (**B**) Densitometry analysis of a minimum of n = 100 crypts per mouse was performed and z-scores generated using the 10-week wild-type mice as the control population. Each dot represents one crypt. The red dashed line is at z= - 3 and the blue dashed line is at z = -6. Crypts were categorised as positive (z ≥ -3), intermediate (-3 > z ≥ -6) or deficient (z < -6). (**C**) Crypts were categorised according to the z-scores and the proportions of crypts in each category are shown. (**D**) Comparison of the proportion of COX-deficient crypts in C5024T mice at 10 weeks and 50 weeks (p=0.4, Mann- Whitney Test).

## Discussion

In this study we have confirmed the loss of m.5024C>T in multiple rapidly dividing tissues of a mouse model of mitochondrial disease and have demonstrated that mutation loss occurs in tissues with the highest rate of cell division, and that post-mitotic tissues, such as skeletal muscle and heart, maintain a steady load of m.5024C>T over time. In addition, we have demonstrated that the mutation loss could be due to a lethal upper limit of the m.5024C>T mutation load in stem cell populations with cell-cycle linked stringent mtDNA replication.

Different dynamics of mtDNA mutations in mitotic and post-mitotic tissues were initially revealed in patients with Pearson syndrome. These patients begin with severe anaemia due to the pathogenicity of large-scale mtDNA deletions, but then lose the deletion in blood rapidly in a few years [32–34] and recover from the anaemia. However, a high proportion go on to develop Kearns-Sayre Syndrome which mainly affects post-mitotic tissues [32, 34]. The most common mtDNA point mutation, m.3243A>G, was later reported to be lost in blood over time, and previous studies have shown a significantly lower load of m.3243A>G in epithelial cells compared with smooth muscle taken from the same organ [12, 14, 35, 36]. The development of the C5024T mice, which recapitulates many of the molecular and biochemical features of primary mtDNA disease [37], has allowed further investigation of the mechanisms underlying the loss of mtDNA mutations selectively from dividing tissues. In agreement with previous studies [22, 37], our data confirm the decline of m.5024C>T load in several mitotic tissues with age in these mice. We show a loss of the mutation in the spleen, the epithelium of the stomach and intestine. Such loss is absent in organs with both dividing and non-dividing cells, such as kidney, pancreas, lung and liver, and in post-mitotic tissues such as muscle, heart and brain, which could explain the tissue-specific phenotype in adult patients with mtDNA disease [38]. Both our experimental and *in silico* analyses have provided evidence that the rate of loss of m.5024C>T is directly related to the proliferation rate of the tissue. Small intestinal epithelium, which has a high rate of turnover, lost the mutation more rapidly than gastric epithelium in mutant mice. In addition, within the epithelium of the stomach, the pyloric epithelium, which has a higher turnover rate, lost the mutation faster than the fundic epithelium.

As with patients harbouring the m.3243A>G mutation [35], the C5024T mice show considerable variation in m.5024C>T load in individual monoclonal intestinal crypts at 10 weeks, and this variation increases significantly by 50 weeks. Each mouse intestinal crypt contains ∼7 effective stem cells, however through a dynamic process of balanced stem cell expansion and loss, one stem cell becomes dominant within the niche within a few weeks [27, 30], meaning that the whole crypt is genotypically monoclonal, and the whole crypt turns over every three days [39]. Therefore, loss of m.5024C>T is most likely to be occurring within the stem cell population; however, we are still unsure as to the exact mechanism by which this is happening. We did not see any significant differences in the proportion of proliferating cells or the frequency of apoptotic cells in crypts between age-matched wild-type and C5024T mice. As apoptosis is a rare and rapid event and is very difficult to measure, it cannot be completely ruled out. However, an alternative hypothesis might be that when stem cells reach the upper tolerance limit for m.5024C>T, competition for stem cell niche space is no longer neutral with ‘fitter’ cells able to outcompete those with the high mutation load. Those stem cells with high m.5024C>T load are then pushed out of the niche and lost by differentiation. Cells with lower m.5024C>T load expand within the niche and then go on to populate the crypt resulting in a lower overall load of m.5024C>T in a relatively short time. This has been shown to occur in reverse in intestinal stem cells which acquire an oncogenic mutation that provides them a selective advantage for clonal expansion and niche succession [40].

In contrast to the mutation dynamics in individual intestinal crypts, we did not observe any mutation loss or changes in variation in the mutational load of m.5024C>T in single skeletal muscle fibres of C5024T mice with age. Our *in silico* modelling suggests that the additional mtDNA replication induced by stochastic stem cell division in the intestinal epithelium can explain this difference. One major assumption we made when establishing the model was that post-mitotic skeletal muscle fibres carried the same mtDNA copy number as stem cells, but realistically each muscle fibre can harbour several thousand copies of mtDNA [41]. However, we assumed that there was an effective population size of mtDNA genomes that are actively replicating within fibres, based on previous studies by our group [42].

Somatic mtDNA mutations occur randomly and clonally expand within individual cells in various tissues (e.g., colon, stomach, heart, skeletal muscle and liver) of normal individuals with age. These mtDNA mutations can result in OXPHOS defects, such as COX deficiency and abnormal cellular function [2, 4, 43–47]. Our data show that the germline m.5024C>T mutation is pathogenic, and we detected COX deficiency in the intestinal crypts at 10 weeks of age. Surprisingly, the proportion of crypts with OXPHOS defects did not change with age, despite a decrease in the m.5024C>T mutation load within the tissue, and by 50 weeks the correlation of OXPHOS defects with the m.5024C>T load at 3 weeks was lost. This suggests that there may be accumulation of random somatic mtDNA mutations which are also causing OXPHOS defects in these mice.

An outstanding question that we are still unable to answer is why clonally expanded pathogenic somatic mutations and the resulting OXPHOS-deficient cells are tolerated in dividing cells of aging mice and humans, but inherited pathogenic mtDNA mutations are lost from rapidly dividing cells of young mice? What changes are occurring with age that are allowing OXPHOS deficiency to persist? Further investigations into age-related changes in stem cell quality control mechanisms are required to address this intriguing question.

In summary, we have shown that there is a loss of m.5024C>T from mitotic cells with age and the rate of loss is proportional to the turnover rate of the tissue, however the exact cellular mechanisms are still unknown. Further work is required to investigate the effects of mtDNA mutations on intestinal stem cell niche dynamics.

## Materials and methods

### Mouse model

Mice carrying the germline m.5024C>T in the mitochondrial tRNA^Ala^ gene were produced as previously described [36] and maintained by backcrossing to C57Bl/6NCrl males (functional *NNT* gene, *COX7A2L/SCAFI*^short^ isoform, *CRB1* gene truncation). Animals were group housed with a 12-hour light-dark cycle and fed ad libitum on a standard chow diet (RM-H Low-Phytoestrogen) or an enhanced diet during breeding and weaning (M-Z low-Phytoestrogen) from Ssniff Spezialdietan GmbH. The study was approved by the Landesamt für Natur, Umwelt and Verbraucherschutz Nordrhein-Westfalen and performed in accordance with the Federation of European Laboratory Animal Science Association guidelines, and the Newcastle University Animal Welfare and Ethical review Board (AWERB425). The number of mice and gastric units/intestinal crypts/skeletal muscle fibres used in the study of heteroplasmy and/or OXPHOS function is summarised in Table S1, S2 and S3.

### Tissue preparation

C5024T mice aged ∼10 weeks and ∼50 weeks and age-matched wild-type controls were humanely killed by cervical dislocation. Mouse organs were harvested and halved: one part was snap-frozen in isopentane cooled in liquid nitrogen (frozen tissue), and the other part was fixed in 10% neutral buffered formalin for 24 hours then dehydrated and embedded in paraffin (FFPE tissue). Ear notch biopsies of mutant mice were acquired at the age of ∼3 weeks to determine the initial mutation burden.

### Total DNA extraction from tissue homogenate and individual intestinal crypts/gastric units/ skeletal fibres

DNA was extracted from snap-frozen organs using DNeasy Blood & Tissue Kits (QIAGEN) and following the manufacturer’s standard protocol. In the study of individual intestinal crypts/gastric units/smooth muscle patches/skeletal muscle fibres, respective frozen samples were cryosectioned at 20 µm. Individual crypts/gastric units/smooth muscle patches (∼30000 µm^2^ in size)/skeletal muscle fibres were randomly selected and collected by using a PALM Laser microdissection system (Zeiss). Collected samples were lysed to obtain DNA as previously described [48].

### Pyrosequencing

The target DNA sequence was amplified by a single round of PCR for pyrosequencing. Sequence and programme information for PCR is in Table S4 and Table S5. Heteroplasmy in each sample was quantified, as previously described [48]. Sequence information for pyrosequencing is in Table S5. Wild-type DNA controls were included to ensure the fidelity of the assay with errors ≤ 3%. All samples were run in triplicate with the mean measurements as the result.

### Modelling mitochondrial dynamics in non-dividing and dividing tissues

The model describes a population of *W* wild-type mtDNA molecules and *M* mtDNA mutations within a single cell. The proportion of wild-type mtDNA is defined as 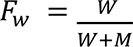 and the proportion of mutant mtDNA (or the mutation load) is defined as 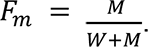. The initial state of the population is *W_0_*, *M_0_*, and the population evolves between time *t = 0* and a specified stopping time, *t = t_stop_*, or until *F*_*m*_ ≥ *L*, where *L* is the upper limit where the mutation is lost. The system evolves according to the following network of reactions and we simulate it using the Gillespie algorithm [49]:

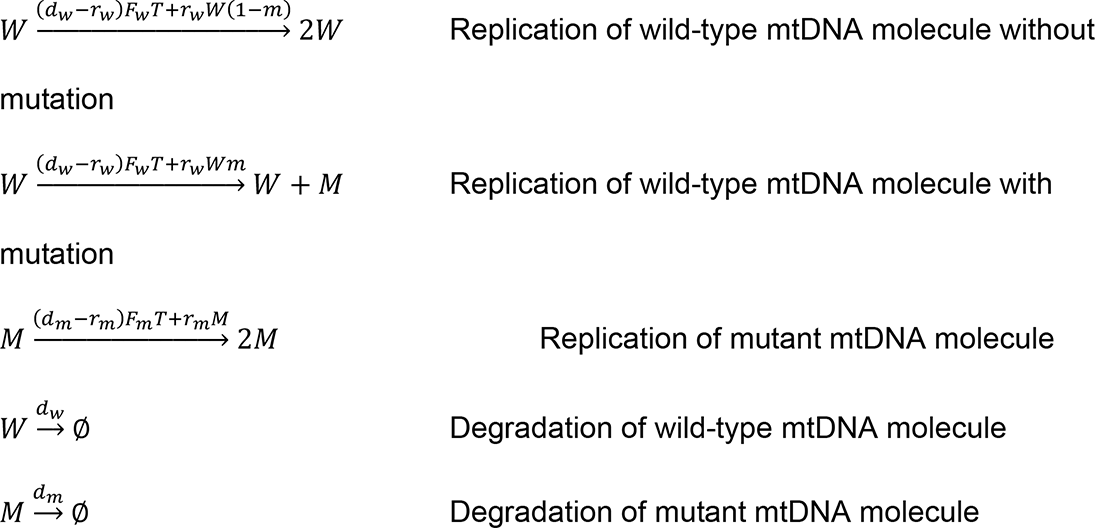

*r_w_* is the rate constant describing replication of wild-type molecules according to first order mass-action kinetics, and *d_w_* is the equivalent degradation rate. *T* is a target, steady-state total mtDNA population size. In the absence of any other known mechanism, copy number homeostasis is modelled here with a linear controller which increases replication when total mtDNA number is lower than *T*, and decreases replication when it is greater than *T*. *m* is the probability that a replication event gives rise to a mutant mtDNA molecule. *r_m_* is the rate constant describing replication of mutant molecules.

To simulate mtDNA population dynamics in a post-mitotic skeletal muscle, we set the model parameters *r_w_*, *d_w_, r_m_, d_m_, m, T* and *L*, chose appropriate initial conditions *W_0_* & *M_0_* and simulated each replication, mutation and degradation event from *t = 0* until *t ≥ t_stop_* (where *t_stop_* is chosen to suit the biological system under examination) using the Gillespie algorithm [49]. To capture the range of possible dynamics from this stochastic model we simulate 1,200 times and examine distributions of e.g. mutation loads, before comparing with experimental results.

#### Stem cell niche model

We also developed an exact, individual-based stochastic model of the dynamics of the stem cell population at the base of epithelial crypts. We define *N*, the number of stem cells in the niche and *N_targ_* the target, steady-state number of stem cells. The initial condition: *N_0_ = N_targ_*. Each cell is defined by the time it was born (*t_b_*), its mtDNA population at birth (*M_b_* & *W_b_*) and the time to next division (*t_div_*). *t_div_* is sampled from the Gaussian distribution with mean *µ* and standard deviation *σ*. We simulate cell divisions by selecting the cell in the niche with the smallest value of *t_div_* and then use the between-division model to simulate mtDNA population from *t_b_* to *t_div_* or until *F*_*m*_ ≥ *L*. To simulate mtDNA replication during cell division, we note that mtDNA copy number increases before cell division [50] and so, if *F*_*m*_ < *L* , we double the mtDNA population before randomly sampling approximately half of the doubled population for each of the two resulting daughter cells. In the absence of mechanistic information about how the niche maintains *N*, we simulate control by assuming that division is always symmetric, giving rise to two stem cells if *N < N_targ_ – δ* and always symmetric giving rise to two non-stem cells if *N > N_targ_ + δ*. Otherwise, the type of the two daughter cells is defined by the parameter *P*, the probability of asymmetric division.

As the mutation load dynamics observed in dissected epithelial crypts is a good proxy for the mutation load dynamics within crypt stem cell niche, we simulated both post-mitotic skeletal muscle fibres (between-division model alone) and mitotic epithelial stem cell niche (stem cell niche model) with the same set of parameters, where appropriate, and parameters specific to the stem cell niche simulation (Table 1). The simulated output visually shows good fitness to the data (Fig. 3). *W_0_* & *M_0_* were estimated from mutation loads experimentally observed at 70 d, assuming initial total copy number was equal to *T*. Post-mitotic skeletal muscle fibres were simulated with *t_stop_ = 730 d*, whereas for the epithelial stem cell niche the *t_stop_* varied for each stem cell division.

### Simulating from models

The output from stochastic models typically requires numerous independent, typically parallel, simulations to capture the distribution of possible outcomes resulting from the random nature of modelled processes. Carrying out this number of simulations requires a lot of CPU time. Models were coded in Julia [53], a scientific programming language, to allow fast simulation and easy deployment across multiple CPUs. Simulation code and experimental data are in a GitHub repository: https://github.com/CnrLwlss/Su_2019.

### Immunofluorescence and image analysis

FFPE tissue was sectioned at 4 µm and placed at 60°C for an hour followed by Histoclear and a descending ethanol gradient. Sections were immersed in 1mM EDTA buffer, pH 8.0 and boiled for 20 minutes in a pressure cooker for antigen retrieval. Specimens were incubated in normal goat serum (10% in 1x TBST, Sigma Aldrich) to reduce non-specific binding of secondary antibodies. Endogenous binding sites of biotin and avidin were blocked using an Avidin/Biotin blocking kit (Vector Laboratories). Sections were incubated in primary antibody cocktails overnight and secondary/tertiary antibodies for two hours at 4°C in a humid chamber with 1x TBST washes in between. Antibodies used were anti-Ki-67 (#12202), anti-Caspase 3, active (cleaved) form (AB3623), Biotin-XX (A10519) and Alexa Fluor secondary/tertiary antibodies (Life Technologies). Regarding nuclei counting, sections were incubated with 0.01 mg/ml Hoechst (Life Technology) for 15 minutes at room temperature in the dark. Sections were mounted using Prolong Gold (Life Technologies).

Sections were imaged using A1+ Confocal Laser Microscope System (Nikon) with an apochromatic lens (x20 magnification, numerical aperture 0.75 and working distance 1.0 mm) providing lasers at four wavelengths (405nm, 488nm, 561nm and 647nm) and NIS-Elements Imaging Software v4.40 (Nikon). Nine-step z-stack images of sections were captured at 1 µm intervals. Individual channels were scanned in sequence. Laser and camera settings were kept identical for all samples. Image processing was conducted using NIS-Elements Imaging Software v4.40 (Nikon). Z- stack images were integrated to acquire maximum intensity projection, i.e. signal intensity at optimal focus at each pixel site. Signals of Ki-67, nuclei and cleaved caspase-3 were manually counted.

### NBTx Histochemistry

NBTx histochemistry was performed on sections of frozen colonic epithelium as previously described [31]. Sections were imaged using the Aperio virtual pathology system (Leica Microsystems, UK) and densitometry analysis performed using Image J [54].

## Acknowledgements

This study was funded by the Newcastle University Centre for Aging and Vitality supported by the BBSRC, EPSRC, ESRC, and MRC as part of the cross-council Lifelong Health and Wellbeing Initiative (DMT, LCG); the Wellcome Centre for Mitochondrial Research (203105/Z/16/Z) (LCG and DMT); UK NIHR Biomedical Research Centre in Age and Age Related Diseases award to the Newcastle upon Tyne Hospitals NHS Foundation (DMT); the United Mitochondrial Disease Foundation (project 13-053R) (JBS); the Max Planck Society (JBS).

## Supplemental Information

**Figure S1.**
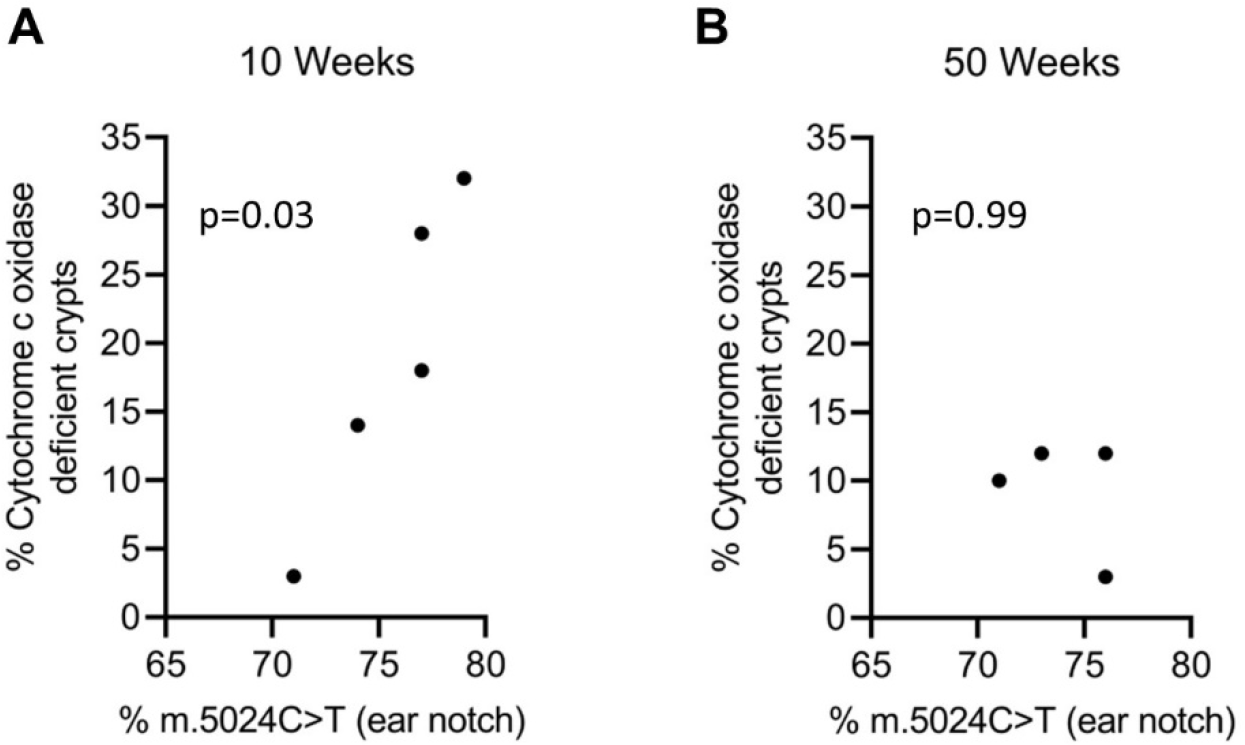
Correlation between the percentage of cytochrome *c* oxidase deficient colonic crypts (measured using NBTx histohemistry) with the mutation load from ear notch at 3 weeks of age. (A) 10-week and (B) 50-week C5024T mice. Spearman’s rank correlation analysis.

**Table S1:** The number (n) of the 10-week and 50-week C5024T mice included in the heteroplasmy analysis for each tissue/organ (Figure 1). Ear notches were collected at weaning (three weeks). SI, small intestine.

| Tissue/organ homogenate | 10-week (n) | 50-week (n) |
| --- | --- | --- |
| Ear notch | 7 | 7 |
| Gastric smooth muscle | 7 | 7 |
| SI smooth muscle | 7 | 6 |
| Heart | 7 | 7 |
| Skeletal Muscle | 7 | 7 |
| Brain | 7 | 7 |
| Kidney | 7 | 7 |
| Pancreas | 6 | 7 |
| Lung | 6 | 7 |
| Liver | 7 | 7 |
| Spleen | 7 | 7 |
| Fundic epithelium | 7 | 6 |
| Pyloric epithelium | 3 | 3 |
| SI epithelium | 7 | 5 |

**Table S2:**
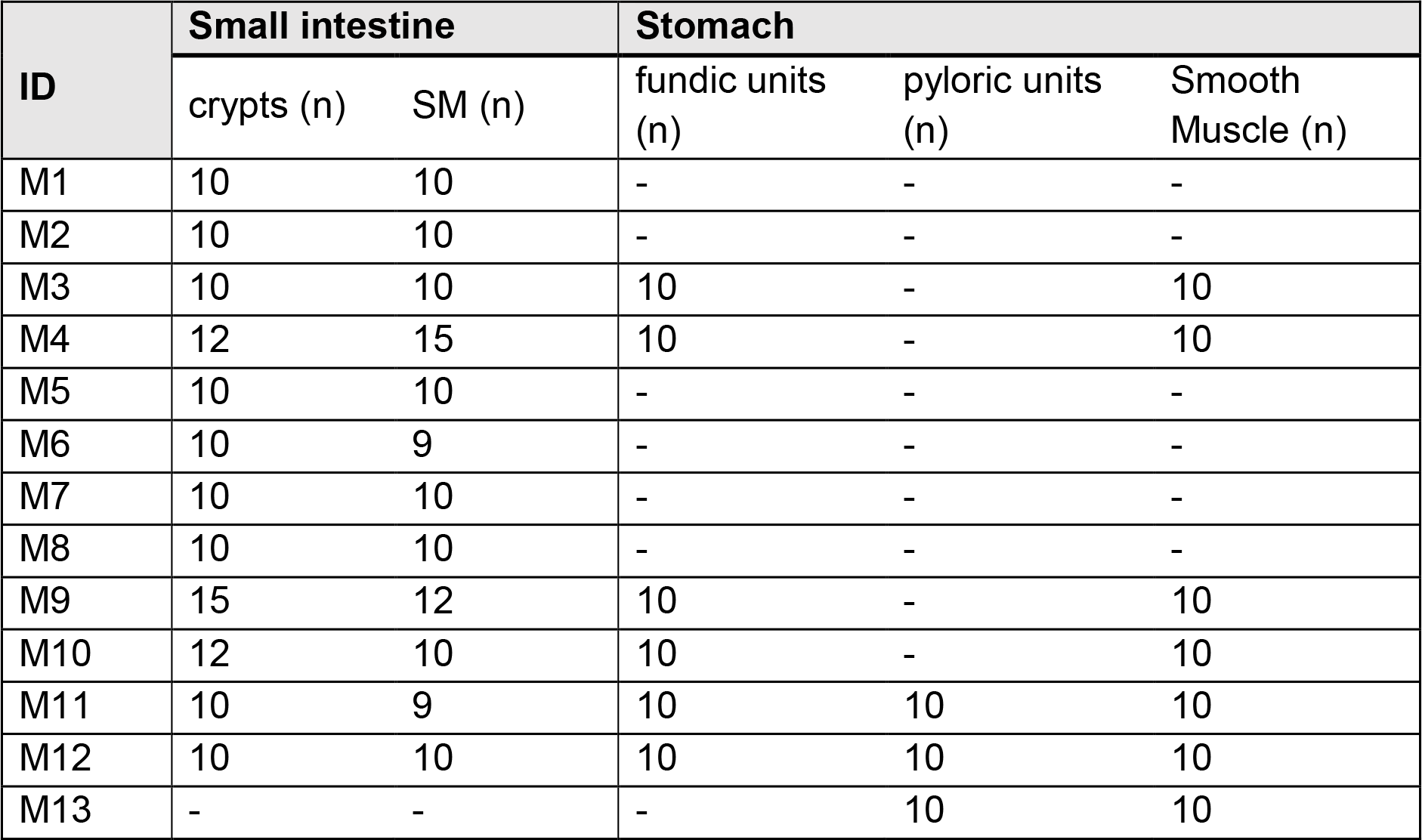
The number (n) of intestinal crypts/gastric units of fundus or pylorus/areas of smooth muscle included for genetic statistical analysis (Figure 2). M1, M2, M5, M6, M7 and M8 were killed at 10 weeks. M3, M4, M9, M10, M11, M12 and M13 were killed at 50 weeks.

**Table S3:** A summary of the Mouse colonic tissue used for NBTx analysis (Figure 5).

| Mouse colonic tissue used for NBTx analysis |  |  |
| --- | --- | --- |
| Mouse ID | m.5024 C>T load (%) | Age (weeks) |
| M5 | 71 | 10 |
| M6 | 74 | 10 |
| M7 | 77 | 10 |
| M8 | 79 | 10 |
| M14 | 77 | 10 |
| WT1 | 0 | 10 |
| WT2 | 0 | 10 |
| WT3 | 0 | 10 |
| M14 | 71 | 50 |
| M15 | 73 | 50 |
| M16 | 76 | 50 |
| M17 | 76 | 50 |
| WT4 | 0 | 50 |
| WT5 | 0 | 50 |
| WT6 | 0 | 50 |
| WT7 | 0 | 50 |
| WT8 | 0 | 50 |
| WT9 | 0 | 50 |

**Table S4:** Thermal cycler programme for amplifying DNA segments spanning mt.5024 mutation site.

| Step | Temperature | Duration | Cycles |
| --- | --- | --- | --- |
| Initial denaturation | 95°C | 10 minutes | 1 |
| Denaturation | 95°C | 30 seconds | 38 |
| Annealing | 62°C | 30 seconds |  |
| Extension | 72°C | 30 seconds |  |
| Final Extension | 72°C | 10 minutes | 1 |
| Hold | 4°C | ∞ |  |

**Table S5:**
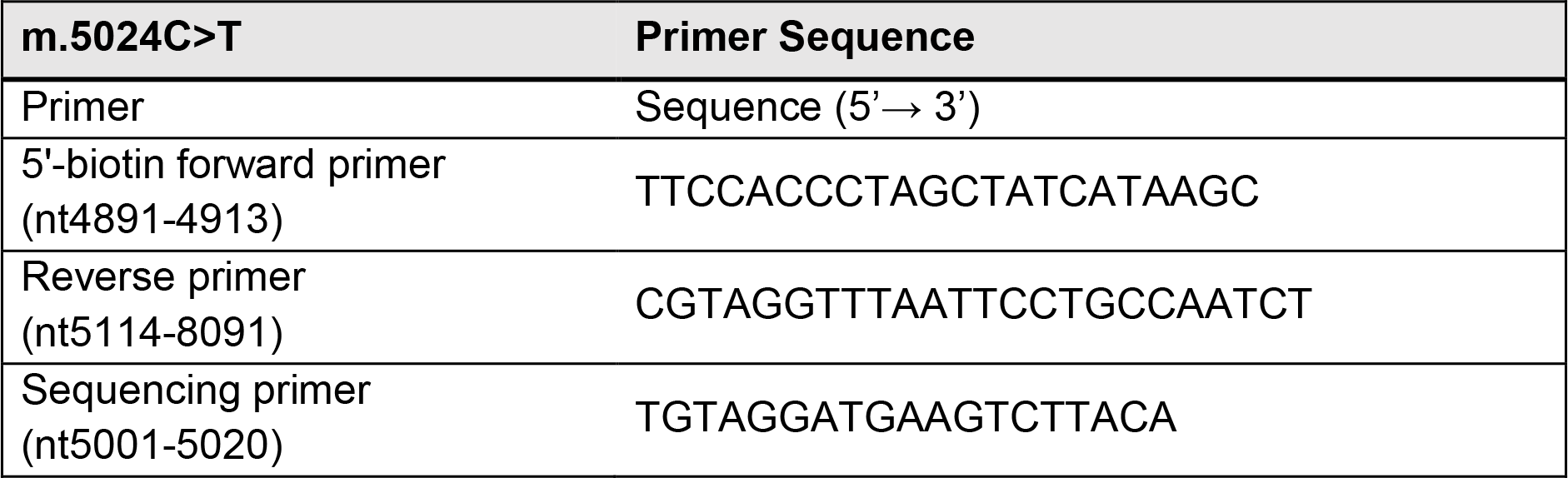
Primer sequences information for pre-pyro PCR and pyrosequencing.

